# The payoffs and tradeoffs of hygienic behavior: A large-scale field study on a local population of honey bees

**DOI:** 10.1101/2021.06.07.447383

**Authors:** Rya Seltzer, Paz Kahanov, Yosef Kamer, Amots Hetzroni, Małgorzata Bieńkowska, Abraham Hefetz, Victoria Soroker

**Affiliations:** Agricultural Research Organization, The Volcani Center, Rishon LeZion, Israel; School of Zoology, George S. Wise Faculty of Life Sciences, Tel Aviv University, Ramat Aviv, Israel; The Mina and Everard Goodman Faculty of Life Sciences, Bar-Ilan University, Ramat Gan, Israel; Research Institute of Horticulture in Skierniewice, Apiculture Division, 24-100 Pulawy, Poland; Faculty of Marine Sciences, Ruppin Academic Center, Mikhmoret, Israel

**Keywords:** *Apis mellifera*, breeding, *Varroa* mite, integrated pest management, honey production

## Abstract

Honey bees (*Apis mellifera*) are exposed to a variety of risk factors, but the ectoparasitic mite *Varroa destructor* and its associated viruses are considered to be the most significant problem worldwide. It has been widely recognized that honey bee stocks resistant to the mites are an essential part of any sustainable long-term management of *Varroa*. The aim of this study was to evaluate the efficacy of hygienic behavior in a local population of honey bees in order to reduce *Varroa* infestation. A bi-directional selection for high and low rates of hygienic behavior was carried out in Israel using either queen artificially inseminated or naturally mated. Colonies were screened for performance: population size, honey production, control of *Varroa* infestation, and the level of hygienic behavior. Furthermore, we examined the costs and benefits of selection using measurements of colony performance. Either way, selected lines should be tested for trade-offs and benefits to ensure their productivity. The selection process revealed that the trait is heritable. Maternal phenotype has a significant effect on *Varroa* load, as colonies founded by hygienic daughter queens showed a significantly lower parasite load. No major trade-offs were found between the rate of hygienic behavior, honey yield, and population size. Measuring the direct benefits of hygienic behavior through colony performance suggests that breeding for this trait makes bees more resistant to *Varroa destructor*. These results are promising for our successful local bee breeding programs in a Mediterranean climate.

## Introduction

The Western honey bee, *Apis mellifera*, is the main pollinator of agricultural crops globally (Delaplane and Mayer 2000; Klein et al. 2007; Dolezal et al. 2016). Over time, honey bee colonies have been selected for commercially important traits such as productivity (i.e., honey yields), colony strength, low swarming, and gentle temperament. High colony losses in the last two decades have raised awareness of declining honey bee health and scientific efforts to determine and mitigate the causes of colony loss have been initiated (VanEngelsdorp and Meixner 2010; www.coloss.org). Although honey bees are exposed to a variety of risk factors, infestation by the ecto-parasitic mite, *Varroa destructor*, is considered to be the most significant health problem of *A. mellifera* worldwide (Genersch et al. 2010; Plettner et al. 2016). The *Varroa* mite is a highly specialized parasite on pupae and adult honey bees, feeding on their fat body (Ramsey et al. 2019). Beside its direct harm to the bees, it transmits about 18 different pathogenic viruses that are practically “injected” into the bees during mite’s feeding. (Rosenkranz et al., 2010). These viruses weaken the bees, and cripple them, as in case of deformed wing virus (DWV) (Francis et al. 2013; Mondet et al. 2014; Zioni et al. 2011), as well as causing immunosuppression (Gregory et al. 2005; Ryabov et al. 2014; Zanni et al. 2017) and learning disabilities (Rosenkranz et al., 2010). The devastating impact of these viruses is further synergized by various agrochemicals (Simon-Delso et al. 2014; Steinhauer et al., 2018; Yang and Cox-Foster 2005).

In order to fight this parasite, beekeepers have used a variety of mite control methods, none of which proved fully satisfactory (Soroker et al., 2018). Moreover, the extensive use of synthetic acaricides, especially in large beekeeping operations, may leave residues that are toxic to bees and the consumers of hive products (Mullin et al. 2010). Furthermore, over time the mites develop resistance to all synthetic acaricides available on the market, which renders its control critical (Sammataro et al. 2005; Rosenkranz et al. 2010). For example in Israel, following years of successful use of Fluvalinate and Coumaphos for mite control, their efficacy diminished and these products are no longer in use (Afik, Ministry of Agriculture Extension Services, personal communication). As an alternative, local Amitraz-based products are currently used, but their efficacy against *Varroa* is declining (Zarchi, Ministry of Agriculture Extension Services, personal communication). This situation necessitates the development of a sustainable strategy for *Varroa* management that integrates a number of approaches. Development of mite resistant honey bee stocks is widely recognized as an essential part of any sustainable integrated *Varroa* management (Dietemann et al. 2012; Sammataro and Avitabile 2011; Spivak and Gilliam 1998).

Social insects, including honey bees, display natural resistance mechanisms against pests and pathogens, which involve both physiological and behavioral traits (Evans and Spivak 2010). While *Varroa* infestation typically leads to colony failure within one to two years (Rosenkranz et al. 2010), some colonies of *Apis mellifera* from different parts of the world survive without being chemically treated (Büchler et al. 2010). Although the mechanisms leading to such resistance are not entirely clear, behavioral traits are likely to play a role in these naturally resistant genotypes (Locke 2016). Several stocks of honey bees have been selectively bred for resistance to *Varroa* by phenotypic selection. The most prominent of these stocks in the USA are the Minnesota Hygienic Bees (Spivak and Euter 2001), the USDA-bred “Russian” bees (Rinderer et al. 2001), and the Varroa-Sensitive Hygiene (VSH) stock that was recently transitioned into the “Pol” line (Danka et al. 2016). Breeding efforts elsewhere include, but are not limited to, breeding program in Canada (Guarna et al. 2015), selection programs in Germany (Gempe et al. 2016), France (Le Conte et al. 2011) as well as in other countries as recently reviewed by Le Conte et al. (2020).

Most of the above programs relied on the improvement of brood-targeted hygienic behavior, the impact of which has been extensively investigated (Leclercq et al. 2017). It has been described as a complex behavior involving the detection, uncapping, and removal of damaged brood (Spivak and Gilliam 1998). In case of *Varroa* infestation, this behavior apparently interferes with the reproduction of the mite (Arathi and Spivak 2001; Zakar et al. 2014). Between-generation, comparisons of hygienic behavior performance demonstrated a significant genetic component for this behavior (Scannapieco et al. 2017;). In addition, several studies reported that the value of hygienic behavior heritability is as high as 0.65 (Boecking et al. 2000; Oxley and Oldroyd 2010).

Research that compared performance of local and imported honey bees indicated that breeding programs should rely on local populations, which are already adapted to the immediate environment. Such local breeding efforts may prevent diseases from spreading among populations while preserving global genetic diversity (Büchler et al. 2014; Meixner et al. 2014; Uzunov et al. 2014). This is a strong argument against the exportation of queens from all over the world. Moreover, practical success of a local breeding program must take into account possible tradeoffs with other commercially desired traits, e.g., productivity, colony strength, and gentle temperament. These may affect acceptance of selected resistant lines by beekeepers (Uzunov et al. 2017; Leclercq et al. 2017). Trade-offs and benefits between traits could be the result of pleiotropy, linkage between traits, or a genetic correlation resulting from the selection on specific individuals which carry several unrelated traits. While beekeepers often advocate importing superior stocks, a recent multinational study showed that local stocks display significant advantages over imported ones (Uzunov et al. 2014; Büchler et al. 2010; Niño and Cameron Jasper 2015, and Uzunov and Brascamp 2017).

More than 50 years ago in Israel, the local honey bee race *A. mellifera syriaca* was actively displaced by *A. mellifera ligustica* that was also mixed over the years with other races mainly, *A. mellifera caucasica* and Buckfast (Soroker et al. 2018). However, we believe that over time the majority of the population had gradually adapted to the local conditions of the region. This environment is characterized by hot dry summers and cold rainy winters with a tendency towards drought years where the colony loss occurs mostly in the summer. *Varroa* infestation further exacerbates summer colony loss in Israel, where a 10-15% loss was recorded in the last decade due to extreme dry and hot weather conditions (V. Soroker, unpublished data). We therefore assume that in the Mediterranean region, social immunity against the *Varroa* mite expressed as hygienic behavior is most crucial when forage is scarce and the population size is in decline.

The aim of this study was to screen the local honey bees in Israel for the level of hygienic behavior and to evaluate its impact on Varroa infestation. While most breeding programs are carried out in Europe and North America (Doke et al. 2015), our experimental program took place in Israel’s Mediterranean climate. In order to quantify the apicultural costs and the benefits of the trait as well as the commercial applicability of selected lines under the conditions of obligatory regular chemical treatment against *Varroa*, we conducted bidirectional selection for high and low hygienic behavior.

## Materials & Methods

The study was conducted at the breeding apiary of the Volcani Center, Agricultural Research Organization (ARO), Israel comprising local bee colonies that had been previously selected for honey yield. The colonies had not received any queens from an alien source since 2008. During 2012-2017, we performed a bi-directional selection program based on queens reared from genetically unrelated colonies that exhibited high or low hygienic phenotype. All selected colonies, regardless of their hygienic performance, had honey production that was above average. Each year, six to ten naturally-mated queens were selected based on their maternal lines (high and low), according to the rate of hygienic behavior and honey production of their colony. Ten to 15 colonies were established for each maternal line. In addition, in 2016 and 2017, artificially inseminated queens with sperm from 8-10 drones from either high or low source colonies were used. The daughters of these queens were naturally mated and used to establish new colonies that were assessed as described below.

The colonies and their subsequent generations in this project were distributed within the apiary area and were assessed for honey yield, hygienic behavior, and colony size. Colonies in which the queen superseded were excluded from analyses. In total, this project included 437 colonies over the years, of which 112 were assessed for *Varroa* infestation (see Table 1). We built a data base containing the pedigree and all the hive assessment data including their maternal phenotype (high or low hygienic behavior) from 2012-2017. To prevent heavy loss, we treated all the colonies twice a year against Varroa mite with Amitraz loaded strips (Galbitraz), in accordance with guidelines from the Ministry of Agriculture Extension Service, first during July-August and the second time during November-December.

**Table 1:** Summary of the types of colonies assessed throughout the study by year, according to their maternal lines and mating techniques.

| Year | Maternal phenotype (number of maternal lines) | Maternal mating techniques | N of colonies tested for honey yield and hygienic behaviour | Seasonal assessments (number per season) | Varroa assessments |
| --- | --- | --- | --- | --- | --- |
| 2012 | general population (14) | naturally-mated | 69 | beginning, middle, and end (once a season) | _____ |
| 2013 | general population (9) | naturally-mated | 50 | beginning, middle, and end (once a season) | _____ |
| 2014 | general population(1), high (4)and low(2) hygienic behavior | naturally-mated | 73 | beginning, middle, and end (three times in the season) | April until end of July (25 colonies) |
| 2015 | general population (1), high (2)and low (2) hygienic behavior | naturally-mated | 98 | beginning, and middle (once a season) | April until end of July (16 colonies) |
| 2016 | high (3) and low (2) hygienic behavior | naturally-mated and artificially inseminated | 69 | beginning, middle, and end (once a season) | April until end of July (37 colonies) |
| 2017 | high (4) and low (1) hygienic behavior | naturally-mated and artificially inseminated | 73 | beginning (once a season) | April until end of July (34 colonies) |

Population size and hygienic behavior were evaluated three times a year according to colony development and seasons. The early season occurs after winter *Varroa* treatment. In our local conditions, this period is typified in by an exponential increase in hive population and nectar flow, and takes place during the end of February until mid-March. The second assessment is referred to as the mid-season, and it occurs just after the spring honey harvest during May and June. The third assessment is referred to as the late season, and it takes place following summer honey extraction and prior to the second *Varroa* treatment during July and August. This period is characterized by a population decline, which remains at a low level until after rain has fallen and some flowering has occurred in October. In 2015, hygienic behavior was evaluated three times in each season, in other words, nine times a year for each colony. This was performed in order to determine the seasonal effect on hygienic behavior. Population size was estimated by measuring the sealed brood area by counting the number of decimeters containing pupae in each of the frames, according to Büchler et al. (2013).

### Hygienic behavior

was measured using the “pin test” involving 100 cells containing red eye pupae (Spivak and Gilliam 1998) as described in detail in Beebook (Büchler et al. 2013). This entailed marking 100 cells and piercing them with an entomological pin #2. The proportion of uncapped and cleaned cells (brood removal) was calculated by comparing pictures taken immediately after pinning, and 24 hours thereafter. The proportions of uncapped and cleaned cells per colony was the basis for the bi-directional selection for high and low hygienic behavior. Selection, however, was based on uncapping behavior only, which was better distributed within the time frame of the test. Consistently, extreme colonies were selected to establish the next generation. Throughout, high hygienic colonies were defined if more than 75% uncapping occurred, and low hygienic colonies were defined if less than _4_5% uncapping occurred, of all pinned brood cells after 24 hours in three independent tests.

### Honey yield

was assessed by weighing honey supers in the spring and summer, while subtracting ten kilograms, the weight of an empty super.

### *Varroa* infestation

was measured weekly, during April and July for four out of the six years of this experiment. It total, we measured infestation in 25 colonies in 2014; 16 colonies in 2015; 37 colonies in 2016; and 34 colonies in 2017, using the method of free-falling *Varroa* on a bottom tray (Dietemann et al. 2013). These colonies were all located in one area of the apiary, and they represented similarly high and low hygienic lines. Hives were placed on a 0.5 × 0.5 cm. screen board floor, and an oiled metal tray was placed under it to record *Varroa* mites that die and/or fall to the bottom within 24 hours. For each colony, the number of mites caught on the trays after 24 hours served as a measure of infestation level. The rate of increase in *Varroa* load over time was estimated and the *Varroa* parasite load (measured as parasite x days) was calculated based on the area under the curve, as a function of time from the first measurement. In particular we calculated the value for each adjacent time points based on the formula for trapeze area (S) calculation, when one of the trapeze bases is *Varroa* number at time t and the other base is its value at t+1,while the height of the trapeze is the time between the measurements in days. We subsequently summed all the S values to calculate the *Varroa* load over the entire period.

### Statistical analysis

was performed on seasonal and annual measurements of several dependent variables for the same colony. These measurements were taken throughout the year and analyzed in a compatible model. We analyzed how the selection process affected hygienic behavior of the progeny using a three-way ANOVA with repeated measurements. For this, we took into consideration the following variables: maternal mating type, maternal phenotype, assessment season, and all their interactions. The variables were: seasonal measurements of hygienic behavior (proportions of uncapping and cleaning) and colony size (sealed brood area). The fixed effects analyzed were mating type, maternal phenotype, assessment season and all their interactions. Random effects were year of testing and colony number nested within mating type and maternal phenotype. Significant main effects were examined by the Tukey HSD test. A two-way ANOVA model was applied for annual measurements of spring, summer, annual honey yield, and *Varroa* infestation. The fixed effects that were analyzed were maternal phenotype, year of testing and their interactions. Pairwise association between hygienic behavior (uncapping and cleaning) and honey yield were tested by Pearson Correlation. Significance was set at alpha=0.05. All statistical tests were carried out using the JMP 14 Statistical Program (SAS, USA).

## Results

We tested the relationship between uncapping and cleaning behaviors. Figure 1 represents the data assembled over the five years of the study. We found a significant and highly positive correlation between the two components of hygienic behavior: uncapping and cleaning (Pearson, r = 0.87, *p<*0.0001). Still, the very high rate of uncapping was not always followed by the high rate of cleaning.

**Fig. 1:**
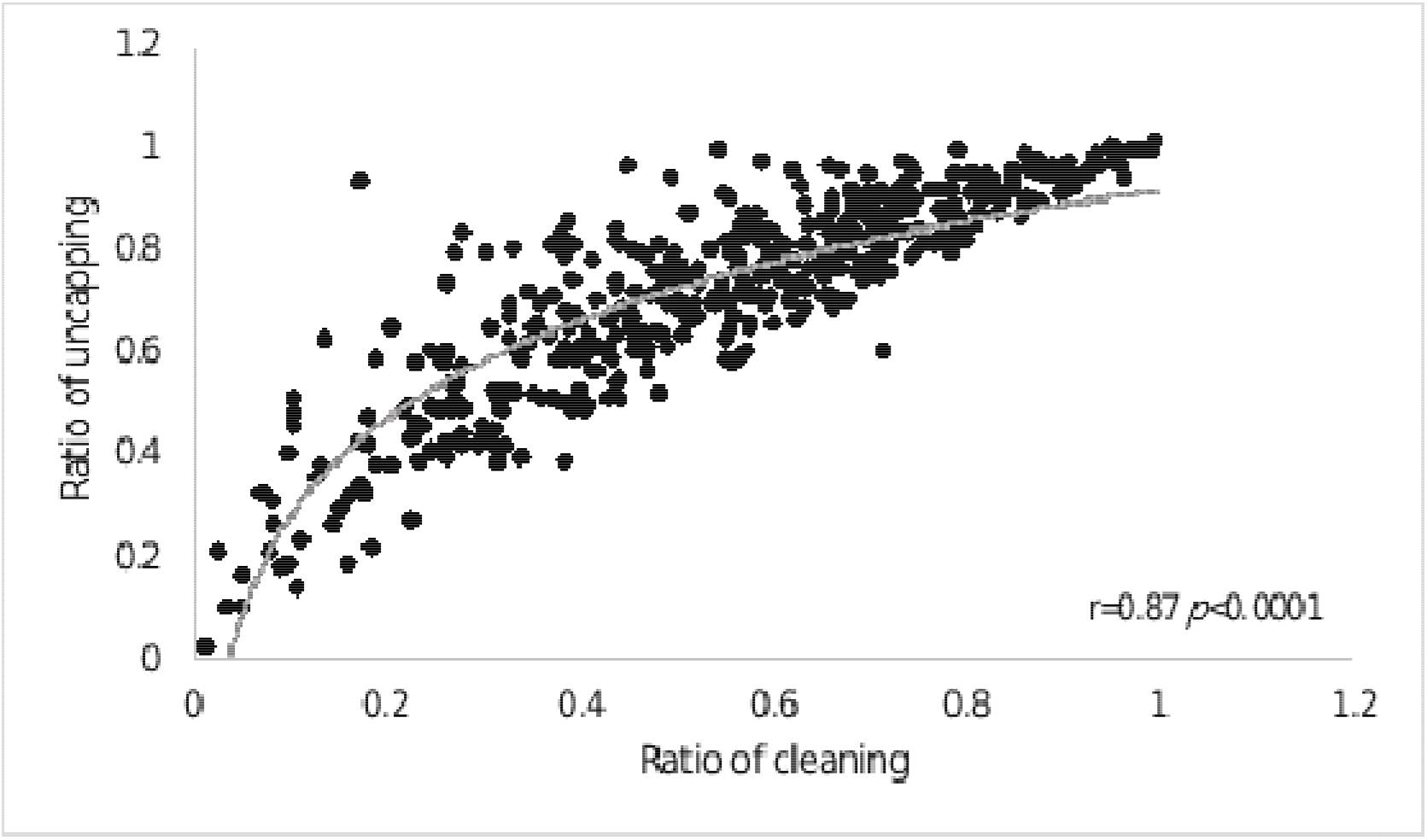
The correlation between the rates of uncapping and cleaning behavior along the tested years. Each dot indicates a hygienic test, and r and *p* are values of Pearson correlation.

Colony size as assessed by sealed-brood area was affected by the maternal phenotype (F_(1,4)_ =4.4, *p* =0.03). Progeny of low hygienic colonies had more sealed brood on average (39.4± 5.3 dm (±SE)) than progeny of high hygienic colonies (36.6± 5.2 dm (±SE)). Maternal mating type did not have a significant effect on sealed-brood area (F_(1,14)_ =0.09, *p* =0.75).

However, as expected, there was a significant effect of seasonality on colony population size (F_(2,662)_=559, *p*<0.0001, Fig.). In early season the average sealed brood area was 46 ± 3.6 dm (± SE); in mid-season it was almost the same with an average of 47.2 ± 3.7 dm (± SE); in late season, however, it dropped significantly to an average 24.5 ± 3.7 dm (± SE). Despite these seasonal fluctuations in population size, we found no significant effect of seasonality on uncapping: (F_(2,836)_ =1, *p*=0.34, Fig._2_). Cleaning behavior was significantly higher in mid-season (0.57 ± 0.03 (± SE)) compared to late season (0.52± 0.03 (± SE)) (F_(2,836)_ =1.05, *p*=0.01, Fig. 2).

**Fig. 2:**
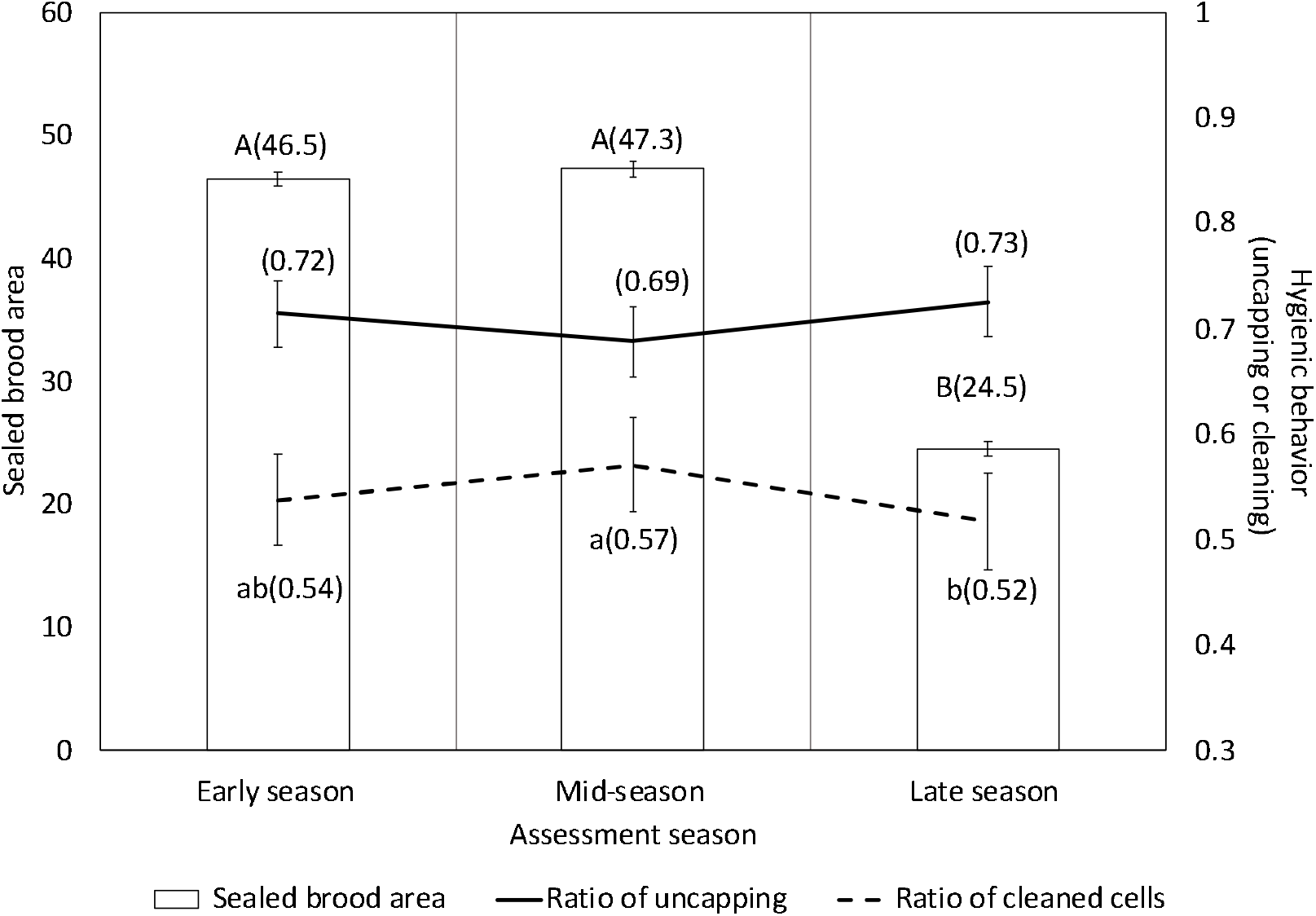
Seasonal changes in sealed brood area and in rate of hygienic behavior. The bars represent sealed brood area (average ±SE) and lines represent hygienic behavior: proportion of uncapping (solid line) or cleaning (dashed line) at different times of the year. The values of average rate of seasonal performances are presented in parentheses. The x-axis represents time of assessment relative to the honeybee season: early season, mid-season, and late season. Significant differences between the seasonal measurements (population size and hygienic behavior) are labeled by different letters (post-hoc Tukey’s HSD, *p*<0.05). The average seasonal performance is presented in parentheses.

The selection according to the maternal phenotype was successful (Table 2). Throughout the duration of the research, the high hygienic progeny colonies had significantly higher levels of both uncapping and cleaning behaviors compared to the low hygienic progeny colonies (high and low respectively, for uncapping 0.8±0.01 vs. 0.6±0.01 (mean ±SE) *p*<0.0001, and for cleaning 0.6±0.03 and 0.4±0.03 (mean ±SE) *p*=<0.0001). The maternal mating type had a significant effect on the cleaning behavior, but not on the uncapping behavior (Table 2). In progeny that were tested for two generations, there was a significant interaction between the maternal phenotype and mating type for both uncapping and cleaning (Table 2), accentuating the differences between the high and low selection lines. Progeny of high hygienic queens that were artificially inseminated had average uncapping behavior of 0.75±0.04 and average cleaning behavior of 0.59±0.04. By comparison, progeny of low hygienic queens that were artificially inseminated had an average uncapping behavior of 0.4±0.03 and average cleaning behavior of 0.26±0.04. On the other hand, for the progeny of the naturally-mated queens, the difference between the two phenotypes were more moderate. Progeny of the naturally-mated high hygienic queens had an average of uncapping and cleaning behaviors of 0.77±0.04 of 0.62±0.02, respectively. Progeny of naturally-mated low hygienic queens seem to have lost their low parental phenotype and showed an average of 0.70±0.03 and 0.55±0.02 uncapping and cleaning behaviors, respectively. Regarding the random variables, colony identity was the only parameter that was significant in our model (uncapping: F_(261,477)_=2.66, *p<*0.0001 and cleaning: F_(261,477)_=2.2, *p<*0.0001 (Table 2)).

**Table 2:** The effect of maternal phenotype (high or low hygienic behavior), maternal mating type (naturally-mated or artificially inseminated), and the interaction among them on hygienic parameters (cell uncapping and cell cleaning). Significant differences in two-way ANOVA among tested groups are marked by an asterisk (colony identity and the year of measurement were taken into consideration in our model as random effects).

|  |  |
| --- | --- |
|  | Response |

Table 2: The effect of maternal phenotype (high or low hygienic behavior), maternal mating type (naturally-mated or artificially inseminated), and the interaction among them on hygienic parameters (cell uncapping and cell cleaning).
| Variables | Proportion of uncapped cells | Proportion of cleaned cells |
| --- | --- | --- |
| Maternal phenotype | $F_{(1,7)}=35$,<br>$p < 0.0001^*$ | $F_{(1,7)}=36$,<br>$p < 0.0001^*$ |
| Maternal mating type | $F_{(1,7)}=35$,<br>$p = 0.95$ | $F_{(1,16)}=5$,<br>$p = 0.02^*$ |
| Maternal phenotype x Maternal mating type | $F_{(1,7)}=5.8$,<br>$p < 0.0001^*$ | $F_{(1,7)}=8.8$,<br>$p = 0.003^*$ |
| Colony (Random) | $F_{(261,477)}=2.66$,<br>$p = 0.0001^*$ | $F_{(261,477)}=2.2$,<br>$p < 0.0001^*$ |
| Year (Random) | $F_{(3,14)}=1.37$,<br>$p = 0.29$ | $F_{(3,17)}=2.18$,<br>$p = 0.12$ |

### *Varroa* infestation

The parasite seasonal load was assessed using the *Varroa-days* formula, which calculates the level of infestation throughout the tested period. Maternal phenotype had a significant effect on parasite load (F_(1,120)_=123, *p =* 0.00_07_). We found the value of this variable to be significantly lower in high hygienic colonies derived from a high hygienic maternal source (_1305_± _283_) compared to the low hygienic colonies (_2858_ ± _275_). The year of testing had also a significant effect on *Varroa* infestation (F_(,3,120)_=23.4, *p<*0.0001). The extreme fluctuation in *Varroa* infestation was exemplified between the years 2016 and 2015. In 2016, we had the lowest *Varroa* infestation, with an average of _477_ ± _333_ per colony. Conversely, the maximum infestation was measured in 2015, with an average of _5005_± _453_.

The interaction between maternal phenotype and the year was also found to be significant (F_(3,120)_=2.7, *p =* 0.047). We found that in the years with heavy *Varroa* infestation, maternal phenotype had a significant effect (in 2014, F_(1,120)_=11.4, *p =* 0.001 and in 2015, F_(1,120)_=4.6, *p =* 0.033, Fig. 3). Obviously in years with low *Varroa* infestation, the secondary effect of maternal phenotype did not have a significant Impact on the parasite load (in 2016, F_(1,120)_=0.0031, *p =* 0.95 and in 2017, F_(1,120)_=0.9, *p =* 0. 34, Fig. 3).

**Fig 3:**
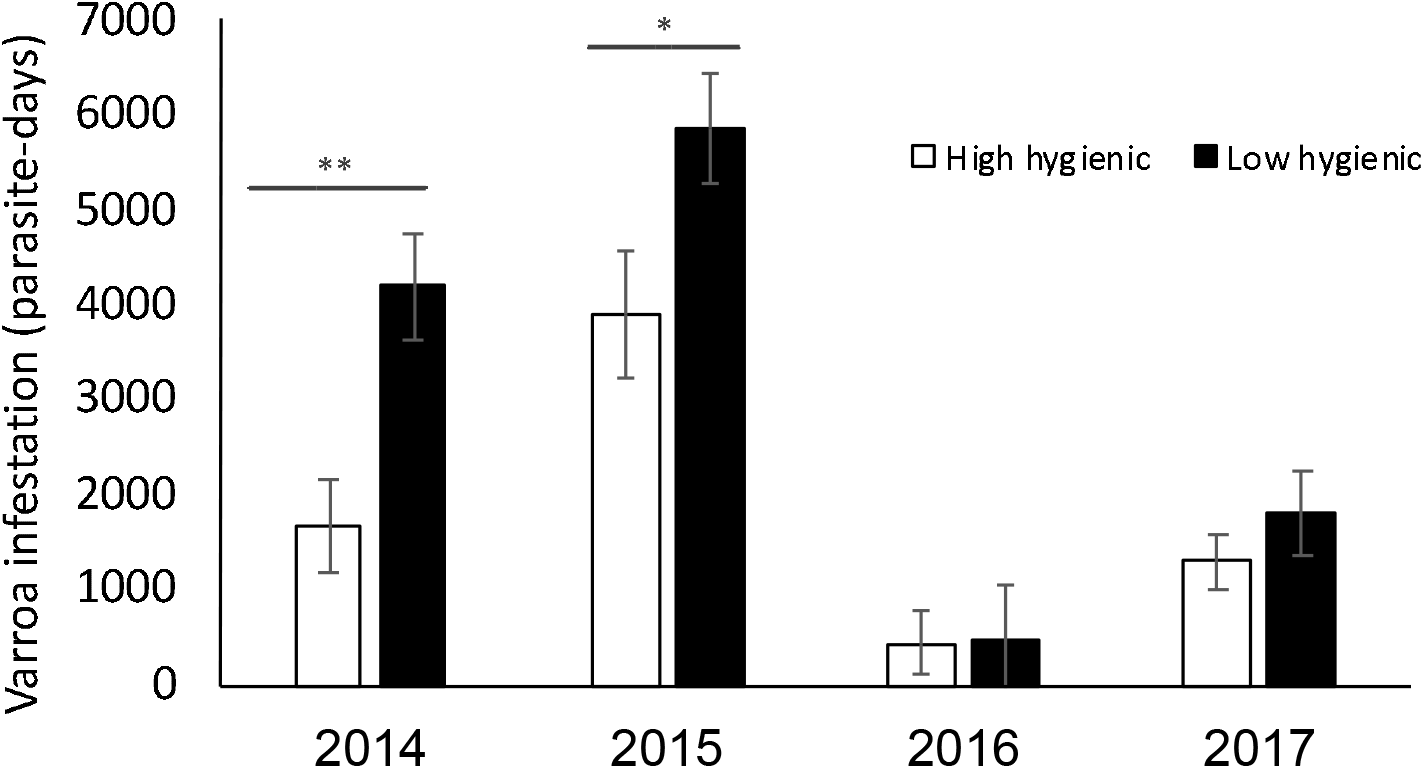
Differences in Varroa infestation between high and low hygienic maternal lines during four years of testing. Varroa infestation is presented in high hygienic (white bar) and low hygienic (black bar) progeny. The data are: average ± SE. The asterisks indicate significant differences in Varroa load between the two groups (post-hoc Tukey’s HSD, *p*<0.05).

### Honey production

A two-way ANOVA analysis of honey yield was performed in order to determine the benefits and identify possible tradeoffs of the selected lines. No significant effect of maternal phenotype was found on honey yields in both seasons (Table 3). In particularly, spring honey yields averaged ± SE: 21.3 ±3.6 kg for high hygienic maternal lines and 21.5 ±3.6 kg for low hygienic lines; summer honey yields averaged: 14.4±2.3 kg for high hygienic maternal lines and 13.8±2.3 kg for low hygienic lines. Annual honey yields averaged 35.2±3.5 kg for high hygienic maternal lines and 34.6±3.5 kg for low hygienic lines. Maternal mating type also had no significant effect on honey yields in both seasons (Table 3). For spring honey yields, artificially inseminated queens’ progeny had on average, _22_±_3_.8 kg while progeny of naturally mated queens had on average 20.8 ±3.5 kg. For summer honey yields, artificially inseminated queens’ progeny had 13.8±2.8 kg while progeny of naturally mated queens had 14.4±2 kg (Table 3). For annual honey yields artificially inseminated queens’ progeny had on average 35.5±4 kg and naturally mated queens’ progeny had 34.3±3 kg (F_(1,11)_=0.2, *p*=0.8 (Table 3). We found that the year of testing (taken into consideration as a random effect) had a significant effect on honey yield. This is a well-known phenomenon which is mainly explained by the differences in environmental conditions between years.

**Table 3:**
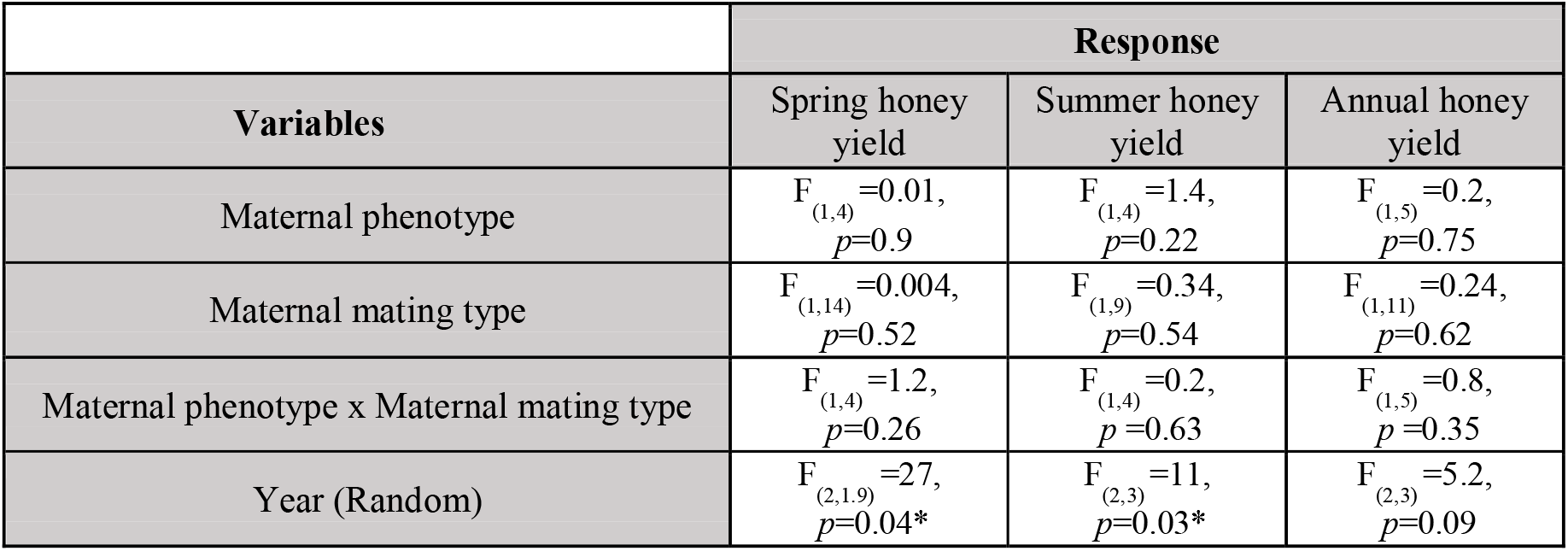
The effects of maternal phenotype type (high or low hygienic behavior) and maternal mating type (naturally-mated or artificially inseminated), and their interaction on honey yields. Significant differences in two-way ANOVA between tested groups are marked with an asterisk. The years of measurement were considered as random effects.

In general, the maternal phenotype, mating technique, and the interaction between them did not have a significant effect on honey yield or on the sealed brood area (Tables 2 and 3). A very low but nonetheless significant negative correlation between uncapping and spring honey yields, was found (Fig. 4A: Pearson, r = -0.137, *p* =0.0093), but practically no correlation regarding cell cleaning (Fig 4B: r = -0.09, *p* =0.07). In contrast, there was a low but positive and significant correlation between summer yields and both measurements of hygienic behavior (Fig. 4C: r = 0.172, *p* =0.003; and Fig. 4D: r = 0.147 *p*=0.0061). Overall, there was no correlation between the annual yield (sum of spring and summer yields) and hygienic behavior (Fig 4E: r = -0.014 *p*=0.78 and Fig. 4F: r = 0.008 *p*=0.87). Taking all of this into consideration, we believe that the correlation was significant, mainly due to a large sample size. This is evident in the pattern of scattered points around the line.

**Fig 4:**
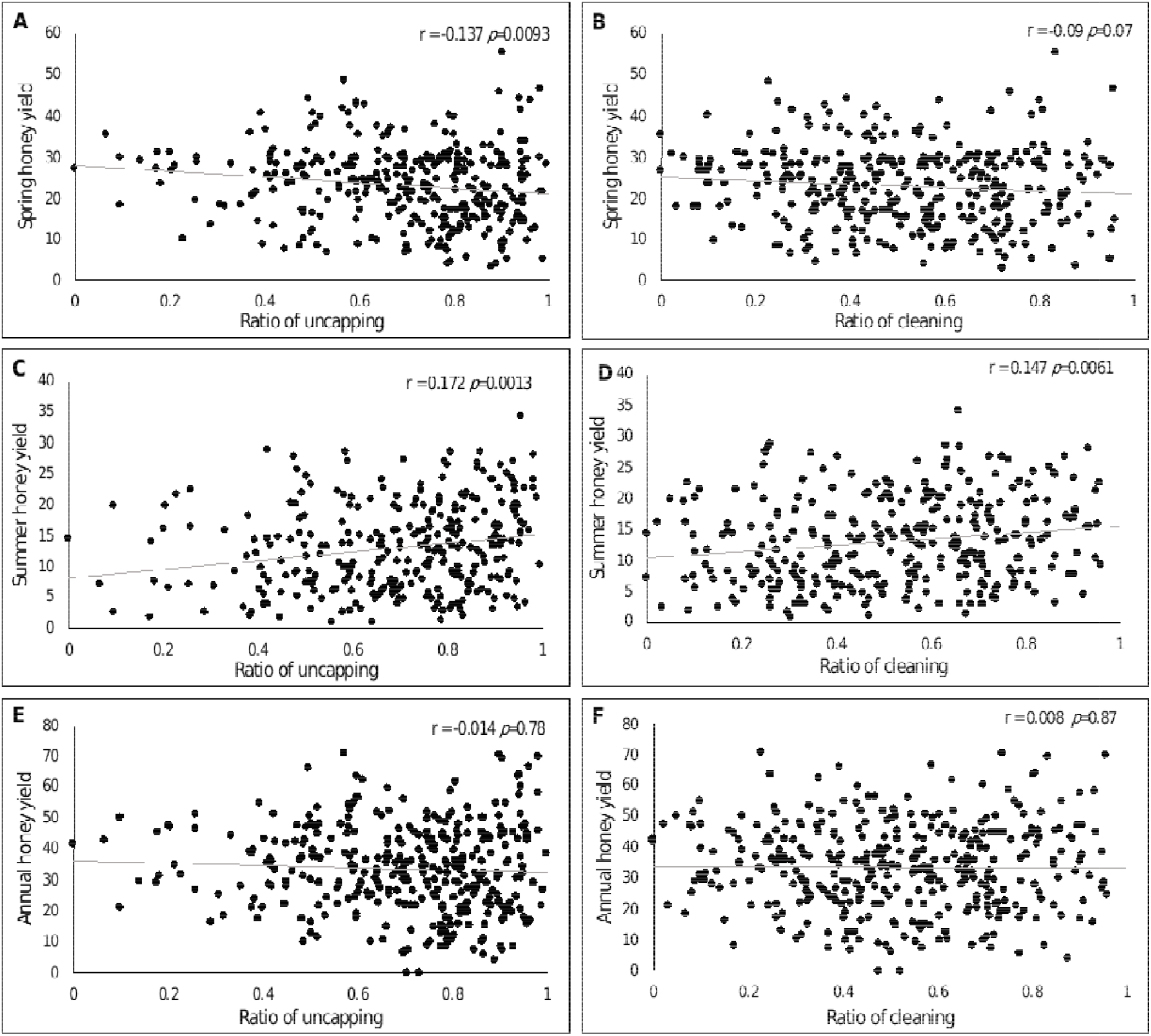
Correlations between uncapping and cleaning of pin damaged brood and colony honey production. Each point represents a different colony. Axis X represents rate of hygienic behavior (proportion of uncapped or cleaned brood cells) and the Y axis represents honey yield in kg. Spring honey yields (A and B), Summer honey yields (C and D) and Annual honey yields (E and F) are shown in the figures. The Pearson correlation is represented by r and *p* values.

## Discussion

Advanced and sustainable management of *Varroa* infestation must include the selection of honeybee lines that show some degree of resistance to the parasite. Here, we evaluated the feasibility of breeding for hygienic behavior as part of *Varroa* management strategy. Our breeding program is unique in that it was carried out in a Mediterranean climate, where colony loss occurs primarily in the summer as opposed to colder regions, where it is most common in winter (Doke et al. 2015). It is well known that there is huge variability in climatic zones and diverse habitats in which honey bees are found. Therefore, we based our hygienic selection on bees that were already adapted to local environmental conditions. This approach supports the preservation of locally-adapted honey bees (Costa et al. 2012; Uzunov et al. 2014). First, we examined the characteristics of hygienic behavior in our study population, namely uncapping and cleaning. As expected, in our study, these hygienic associated behaviors were strongly correlated. Although it is clear that cleaning behavior depends on uncapping, the two together are a prerequisite for successful pest and pathogen control including *Varroa* (Spivak and Danka 2021). Next, we analyzed seasonal effects on hygienic behavior. Seasonality is linked to environmental changes (e.g., temperature, humidity and precipitation), and thus leads to dramatic fluctuations in honey bee population size. The sealed brood area is an indicator of population size, and as a result, it also represents the number of bees at the age of performing hygienic behavior. Some previous studies have suggested an association between the rate of hygienic behavior and seasonal fluctuation in colony size (Uzunov et al. 2014), as well as environmental factors that affect hygienic behavior (Güler and Toy 2013). In contrast, both our results and those of Bigio et al. (2013) demonstrate that hygienic behavior is rather stable along the season and is independent of the population size. We found that in our conditions hygienic behavior is rather stable along the season and fluctuates very little in comparison to the dramatic fluctuations of population size between the seasons.

In agreement with previous research (e.g., Spivak and Reuter 1998; Fly et al. 2014; Zakour and Bienefeld 2014; Danka et al. 2016; de Jesus et al. 2017; Scannapieco et al. 2017), we have found that maternal colony phenotype has a significant effect on hygienic behavior, which emphasizes the potential of breeding for this trait in the local population. We also found a significant interaction between maternal phenotype and mating technique, indicating that artificially inseminated daughters preserve the maternal phenotype better than daughters of naturally-mated daughter queens. This contradicts Bigio et al. (2014a), who claim that there are no advantages in using artificially inseminated queens while breeding for hygienic behavior. Our results indicate that selection based solely on queens is not enough. In fact, the model published by Plate et al. (2019) that simulated the power of selection in a drone controlled set-up, clearly shows that the selection based solely on queens in a large non-selected population is insufficient.

Although pin-killed assays for hygienic behavior is not specific to *Varroa* infected brood (Spivak and Danka 2021) and it is preferable to test such resistance directly by challenging colonies with a parasite, our results clearly showed that lines derived from a high hygienic maternal source, based on pin killed brood assay, also demonstrated a lower parasite load when compared to low hygienic progeny colonies. Moreover, lower loads of *Varroa* mites could result in lower virus infestation (Locke 2012; Kuster et al. 2014; Mondet et al. 2014), which most likely leads to lower virus transmission and improved colony health. Nevertheless, this notion has been questioned by Geffre et al. (2020) as viruses can alter honeybee social behavior. Since this behavior increases contact between the workers and infected brood, the implication of hygienic behavior on viral transmission within and between the colonies as well as the association between *Varroa* infestation, viral load, and social and individual immunity remain to be thoroughly investigated.

Lastly, the feasibility of implementation of honey bees’ hygienic lines in a commercial apiary is tightly linked to the apicultural costs maintaining such lines and whether selection for hygienic trait compromises other desired traits. Seeley (1985) raised a concern about a high cost for hygienic behavior due to the inadvertent removal of healthy brood from the colony. In fact, in our study we found that progeny of low hygienic colonies had more sealed brood, but this could suggest that colonies kept the unhealthy brood rather than the hygienic colonies unintentionally removed the healthy brood. Unfortunately, in our experiments we have not compared the brood quality between the genotypes to test this hypothesis and it should be tested in the future studies. Anyhow, several studies have already shown that hygienic behavior is specifically directed towards damaged brood (Bigio et al. 2014b; Mondet et al. 2016). Potential payoffs and tradeoffs of hygienic behavior with respect to honey yield, propolis production, royal jelly, aggressive behavior, and swarming tendency were reviewed in Leclercq et al. (2017). They concluded that there were no major tradeoffs associated with hygienic behavior. Yet every local breeding program should test the payoffs and tradeoffs of their selected lines, since the latter could carry undesired additional traits. Our analyses of selected lines revealed no impact on annual honey yield or population size except for a small significant negative correlation with uncapping behavior and spring honey yield. There may, however, be a benefit to the trait, supported by the positive correlation between summer honey yield and hygienic performance. The advantage of high hygienic lines was reflected in the strength of the colony during the summer peak of Varroa infestation. We hypothesize that this trend would be of great importance due to the ever-growing abundance of acaricide resistant mites in intensive commercial beekeeping.

In conclusion, our results show that not only the hygienic trait exists in a local population bred for years for honey production, but that selection for this trait reduced *Varroa* infestation without negative impacts on colony size and honey production. Therefore, it is safe to recommend its introduction into local breeding programs as a basis for future integrated *Varroa* management. Moreover, since the literature demonstrates that hygienic behavior is efficient against several bee diseases, such as American foulbrood and chalkbrood (Spivak and Reuter 2001; Leclercq et al. 2017), it will be interesting to test the impact of our selection program on the management of these two diseases in a Mediterranean climate, as well as on the spread of other bee viral diseases.

## Authors’ Contribution

RS, AHefetz and VS designed the experiments. RS, PK, YK, and VS performed the experiments. MB performed and instructed the process of artificial insemination. A Hetzroni constructed the database, and RS analyzed the data. RS, MB, AHefetz and VS wrote the manuscript. All authors agreed to the final version of publication and its content.

## Acknowledgments

We wish to thank the beekeeper Ilia Zaidman for technical assistance with colonies, Dr. Hillary Voet for her help with statistical analysis, Dr. Beatrice Nganso for her critical review on the draft of this manuscript and BARD Binational Foundation grant IS-5078-18 to VS for funding final stages of this research.

